# MRI- and histologically derived neuroanatomical atlas of the *Ambystoma mexicanum* (axolotl)

**DOI:** 10.1101/2020.12.09.418566

**Authors:** Iván Lazcano, Abraham Cisneros-Mejorado, Luis Concha, Juan José Ortíz Retana, Eduardo A. Garza-Villarreal, Aurea Orozco

**Author notes:** **Corresponding authors:** Ivan Lazcano, Instituto de Neurobiología, Laboratorio D-03, Universidad Nacional Autonoma de Mexico (UNAM) campus Juriquilla, Boulevard Juriquilla 3001, Santiago de Querétaro, Querétaro, México, C.P. 76230, Eduardo A. Garza-Villarreal, MD, PhD, Instituto de Neurobiología, Laboratorio B-03, Universidad Nacional Autonoma de Mexico (UNAM) campus Juriquilla, Boulevard Juriquilla 3001, Santiago de Querétaro, Querétaro, México, C.P. 76230.

## Abstract

Amphibians are an important vertebrate model system to understand anatomy, genetics and physiology. Importantly, the brain and spinal cord of adult urodels (salamanders) have an incredible regeneration capacity, contrary to anurans (frogs) and the rest of adult vertebrates. Among these amphibians, the axolotl (*Ambystoma mexicanum*) has gained most attention because of the surge in the understanding of CNS regeneration and the recent sequencing of its whole genome. However, a complete comprehension of the brain anatomy is not available. In the present study we created a magnetic resonance imaging atlas of the *in vivo* neuroanatomy of the juvenile axolotl brain. This is the first MRI atlas for this species and includes 3 levels: 1) 80 regions of interest (ROIs); 2) a division of the brain according to the embryological origin of the neural tube, and 3) left and right hemispheres. Additionally, we localized the myelin rich regions of the juvenile brain. The atlas, the template that the atlas was derived from, and a masking file, can be found on Zenodo at DOI: 10.5281/zenodo.4311937. This MRI brain atlas aims to be an important tool for future research of the axolotl brain and that of other amphibians.

## Introduction

The Mexican axolotl *Ambystoma mexicanum*, (from the Nahuatl, “water monster”) is a mythic native neotenic salamander that has amazed scientists initially for its peculiar life cycle ^1^ [1] and later on for its tissue regeneration capacity ^2^, a trait lost later during vertebrate evolution. Specially interesting is the capacity of the axolotl to regenerate nervous tissue, which has been described for the pallium ^3,4^, the homologue of the cerebral cortex in mammals ^5^, as well as for the spinal cord, both of which are able to regenerate after injury ^6^. Although this capacity has placed axolotl as an animal model for neural regeneration studies, a thorough comprehension of the nervous system of this taxon remains obscure. In this context, and although some axolotl brain structures have been well described by several authors ^5–7^, only an histological atlas of the compete brain of this species is available (https://msu.edu/course/zol/402/atlas/). In general terms, the juvenile-adult brain of axolotl (and other amphibians) is composed by regions similar to those of other vertebrates: olfactory bulb, telencephalon, diencephalon, mesencephalon, and rhombencephalon. Histological procedures provide excellent differentiation of tissue types, albeit having limited sampling of the three-dimensional spatial domain, and the introduction of deformations due to tissue handling. Magnetic resonance imaging (MRI) is a tool that allows the non-invasive assessment of the neuroanatomy of virtually any species that fits the scanner bore. MRI has been employed to generate *in vivo* atlases of the central nervous system (CNS) from different species, *i.e.*, human- ^8^, rodent- ^9–11^, avian- ^12,13^ as well fish ^14,15^, with the advantage of preserving the shape of its parenchyma, as well as vascular and ventricular systems. With this in mind, and to further understand the species anatomy and function, we developed a MRI-based *in vivo* atlas of the juvenile *Ambystoma mexicanum*, and correlated this atlas with a histological visualization of myelin. Given the importance of axolotl as a model species and the fact that it has been brought to the danger of extinction, the creation of this first MRI brain atlas will be an important reference not only for anatomists, but also for biologists, environmentalists or other researchers interested in this specie and other amphibians.

## Results and Discussion

Although the brain of different vertebrates seems structurally different in morphology, it can be subdivided in the same way according to the embryonic origin of the neural tube. In particular, the amphibian brain has been studied in anatomical detail earlier and has been subdivided in olfactory bulb, telencephalon, diencephalon, mesencephalon, and rhombencephalon ^16^. In the present work, we created an average template from 14 juvenile axolotls with a final voxel resolution of 0.040 × 0.040 × 0.040 mm. We manually delineated 80 regions of interest (40 per hemisphere), and the average volume of each region as well as its variability are summarized in **Table 1**. Whole brain volume per hemisphere (left: 13.4 1.66; right: 13.2 1.65) is shown in **Supplementary Figure 1**. We manually segmented these regions based on previous annotations from axolotl histological studies in which techniques such as cresyl violet staining, immunohistochemistry or neuronal tracing were used **Supplementary Table 1**. At the end of segmentation, we are able to create for the first time a MRI atlas of the axolotl brain (**Figure 1**) in which subdivisions of the main structures present in other vertebrates can be observed (olfactory bulb, telencephalon, etc). Moreover, we also segmented the pituitary gland, an endocrine organ which interacts directly and indirectly with the brain.

**Table 1.**
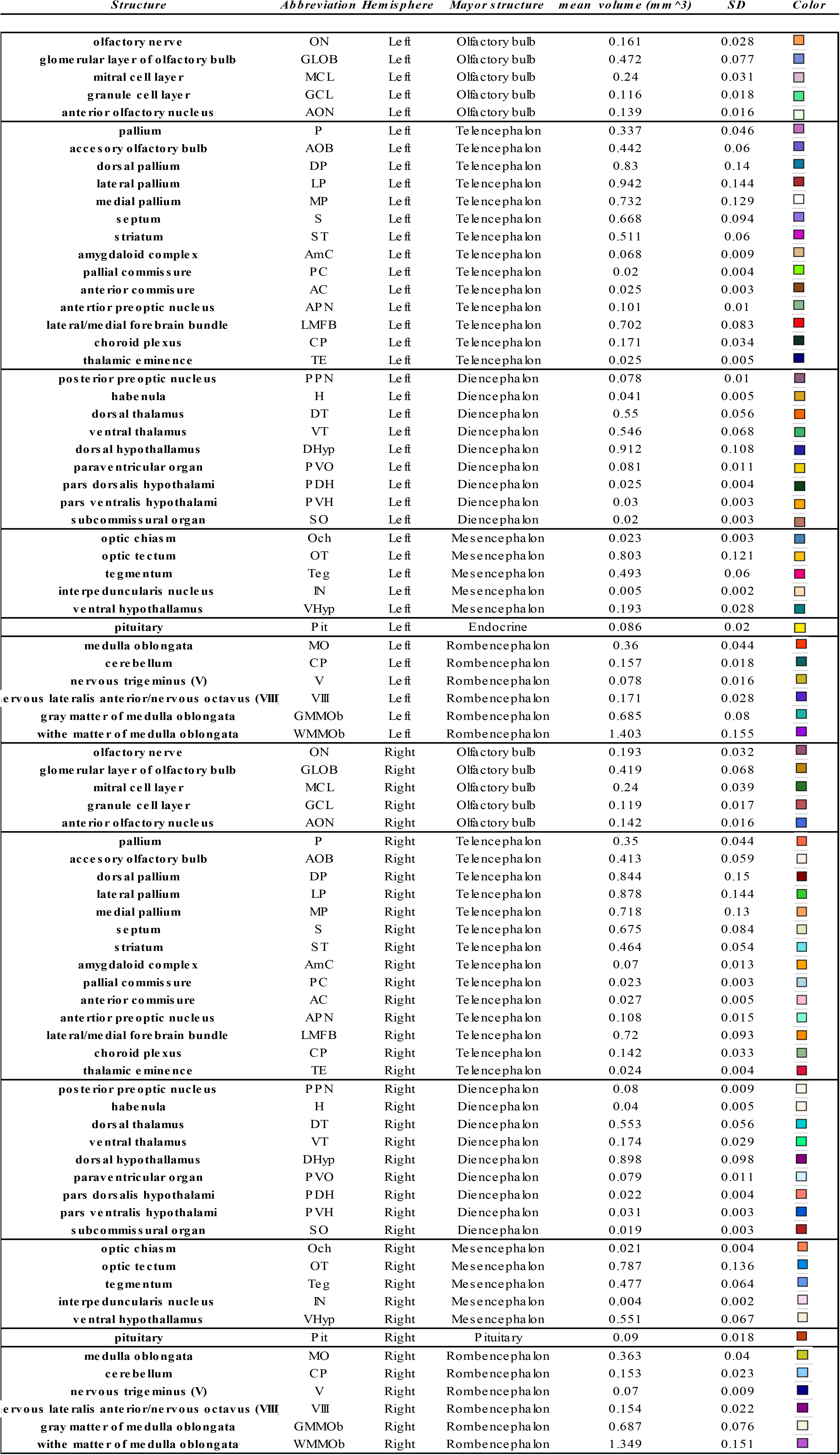
List of ROIs, abbreviations, hemisphere, mayor structure, volume, standar deviation (SD) and color

| *Structure* | *Abbreviation* | *Hemisphere* | *Mayor structure* | *mean volume (mm ^3)* | *SD* | *Color* |
| --- | --- | --- | --- | --- | --- | --- |
| olfactory nerve | ON | Left | Olfactory bulb | 0.161 | 0.028 |  |
| glomerular layer of olfactory bulb | GLOB | Left | Olfactory bulb | 0.472 | 0.077 |  |
| mitral cell layer | MCL | Left | Olfactory bulb | 0.24 | 0.031 |  |
| granule cell layer | GCL | Left | Olfactory bulb | 0.116 | 0.018 |  |
| anterior olfactory nucleus | AON | Left | Olfactory bulb | 0.139 | 0.016 |  |
| pallium | P | Left | Telencephalon | 0.337 | 0.046 |  |
| accessory olfactory bulb | AOB | Left | Telencephalon | 0.442 | 0.06 |  |
| dorsal pallium | DP | Left | Telencephalon | 0.83 | 0.14 |  |
| lateral pallium | LP | Left | Telencephalon | 0.942 | 0.144 |  |
| medial pallium | MP | Left | Telencephalon | 0.732 | 0.129 |  |
| septum | S | Left | Telencephalon | 0.668 | 0.094 |  |
| striatum | ST | Left | Telencephalon | 0.511 | 0.06 |  |
| amygdaloid complex | AmC | Left | Telencephalon | 0.068 | 0.009 |  |
| pallial commissure | PC | Left | Telencephalon | 0.02 | 0.004 |  |
| anterior commissure | AC | Left | Telencephalon | 0.025 | 0.003 |  |
| anterior preoptic nucleus | APN | Left | Telencephalon | 0.101 | 0.01 |  |
| lateral/medial fore brain bundle | LMFB | Left | Telencephalon | 0.702 | 0.083 |  |
| choroid plexus | CP | Left | Telencephalon | 0.171 | 0.034 |  |
| thalamic eminence | TE | Left | Telencephalon | 0.025 | 0.005 |  |
| posterior preoptic nucleus | PPN | Left | Diencephalon | 0.078 | 0.01 |  |
| habenula | H | Left | Diencephalon | 0.041 | 0.005 |  |
| dorsal thalamus | DT | Left | Diencephalon | 0.55 | 0.056 |  |
| ventral thalamus | VT | Left | Diencephalon | 0.546 | 0.068 |  |
| dorsal hypothalamus | DHyp | Left | Diencephalon | 0.912 | 0.108 |  |
| paraventricular organ | PVO | Left | Diencephalon | 0.081 | 0.011 |  |
| pars dorsalis hypothalami | PDH | Left | Diencephalon | 0.025 | 0.004 |  |
| pars ventralis hypothalami | PVH | Left | Diencephalon | 0.03 | 0.003 |  |
| subcommissural organ | SO | Left | Diencephalon | 0.02 | 0.003 |  |
| optic chiasm | Och | Left | Mesencephalon | 0.023 | 0.003 |  |
| optic tectum | OT | Left | Mesencephalon | 0.803 | 0.121 |  |
| tegmentum | Teg | Left | Mesencephalon | 0.493 | 0.06 |  |
| interpeduncularis nucleus | IN | Left | Mesencephalon | 0.005 | 0.002 |  |
| ventral hypothalamus | VHyp | Left | Mesencephalon | 0.193 | 0.028 |  |
| pituitary | Pit | Left | Endocrine | 0.086 | 0.02 |  |
| medulla oblongata | MO | Left | Rombencephalon | 0.36 | 0.044 |  |
| cerebellum | CP | Left | Rombencephalon | 0.157 | 0.018 |  |
| nervous trigeminus (V) | V | Left | Rombencephalon | 0.078 | 0.016 |  |
| nervous lateralis anterior/nervous octavus (VIII) | VIII | Left | Rombencephalon | 0.171 | 0.028 |  |
| gray matter of medulla oblongata | GMMOb | Left | Rombencephalon | 0.685 | 0.08 |  |
| white matter of medulla oblongata | WMMOb | Left | Rombencephalon | 1.403 | 0.155 |  |
| olfactory nerve | ON | Right | Olfactory bulb | 0.193 | 0.032 |  |
| glomerular layer of olfactory bulb | GLOB | Right | Olfactory bulb | 0.419 | 0.068 |  |
| mitral cell layer | MCL | Right | Olfactory bulb | 0.24 | 0.039 |  |
| granule cell layer | GCL | Right | Olfactory bulb | 0.119 | 0.017 |  |
| anterior olfactory nucleus | AON | Right | Olfactory bulb | 0.142 | 0.016 |  |
| pallium | P | Right | Telencephalon | 0.35 | 0.044 |  |
| accessory olfactory bulb | AOB | Right | Telencephalon | 0.413 | 0.059 |  |
| dorsal pallium | DP | Right | Telencephalon | 0.844 | 0.15 |  |
| lateral pallium | LP | Right | Telencephalon | 0.878 | 0.144 |  |
| medial pallium | MP | Right | Telencephalon | 0.718 | 0.13 |  |
| septum | S | Right | Telencephalon | 0.675 | 0.084 |  |
| striatum | ST | Right | Telencephalon | 0.464 | 0.054 |  |
| amygdaloid complex | AmC | Right | Telencephalon | 0.07 | 0.013 |  |
| pallial commissure | PC | Right | Telencephalon | 0.023 | 0.003 |  |
| anterior commissure | AC | Right | Telencephalon | 0.027 | 0.005 |  |
| anterior preoptic nucleus | APN | Right | Telencephalon | 0.108 | 0.015 |  |
| lateral/medial fore brain bundle | LMFB | Right | Telencephalon | 0.72 | 0.093 |  |
| choroid plexus | CP | Right | Telencephalon | 0.142 | 0.033 |  |
| thalamic eminence | TE | Right | Telencephalon | 0.024 | 0.004 |  |
| posterior preoptic nucleus | PPN | Right | Diencephalon | 0.08 | 0.009 |  |
| habenula | H | Right | Diencephalon | 0.04 | 0.005 |  |
| dorsal thalamus | DT | Right | Diencephalon | 0.553 | 0.056 |  |
| ventral thalamus | VT | Right | Diencephalon | 0.174 | 0.029 |  |
| dorsal hypothalamus | DHyp | Right | Diencephalon | 0.898 | 0.098 |  |
| paraventricular organ | PVO | Right | Diencephalon | 0.079 | 0.011 |  |
| pars dorsalis hypothalami | PDH | Right | Diencephalon | 0.022 | 0.004 |  |
| pars ventralis hypothalami | PVH | Right | Diencephalon | 0.031 | 0.003 |  |
| subcommissural organ | SO | Right | Diencephalon | 0.019 | 0.003 |  |
| optic chiasm | Och | Right | Mesencephalon | 0.021 | 0.004 |  |
| optic tectum | OT | Right | Mesencephalon | 0.787 | 0.136 |  |
| tegmentum | Teg | Right | Mesencephalon | 0.477 | 0.064 |  |
| interpeduncularis nucleus | IN | Right | Mesencephalon | 0.004 | 0.002 |  |
| ventral hypothalamus | VHyp | Right | Mesencephalon | 0.551 | 0.067 |  |
| pituitary | Pit | Right | Pituitary | 0.09 | 0.018 |  |
| medulla oblongata | MO | Right | Rombencephalon | 0.363 | 0.04 |  |
| cerebellum | CP | Right | Rombencephalon | 0.153 | 0.023 |  |
| nervous trigeminus (V) | V | Right | Rombencephalon | 0.07 | 0.009 |  |
| nervous lateralis anterior/nervous octavus (VIII) | VIII | Right | Rombencephalon | 0.154 | 0.022 |  |
| gray matter of medulla oblongata | GMMOb | Right | Rombencephalon | 0.687 | 0.076 |  |
| white matter of medulla oblongata | WMMOb | Right | Rombencephalon | 1.349 | 0.151 |  |

**Figure 1.**
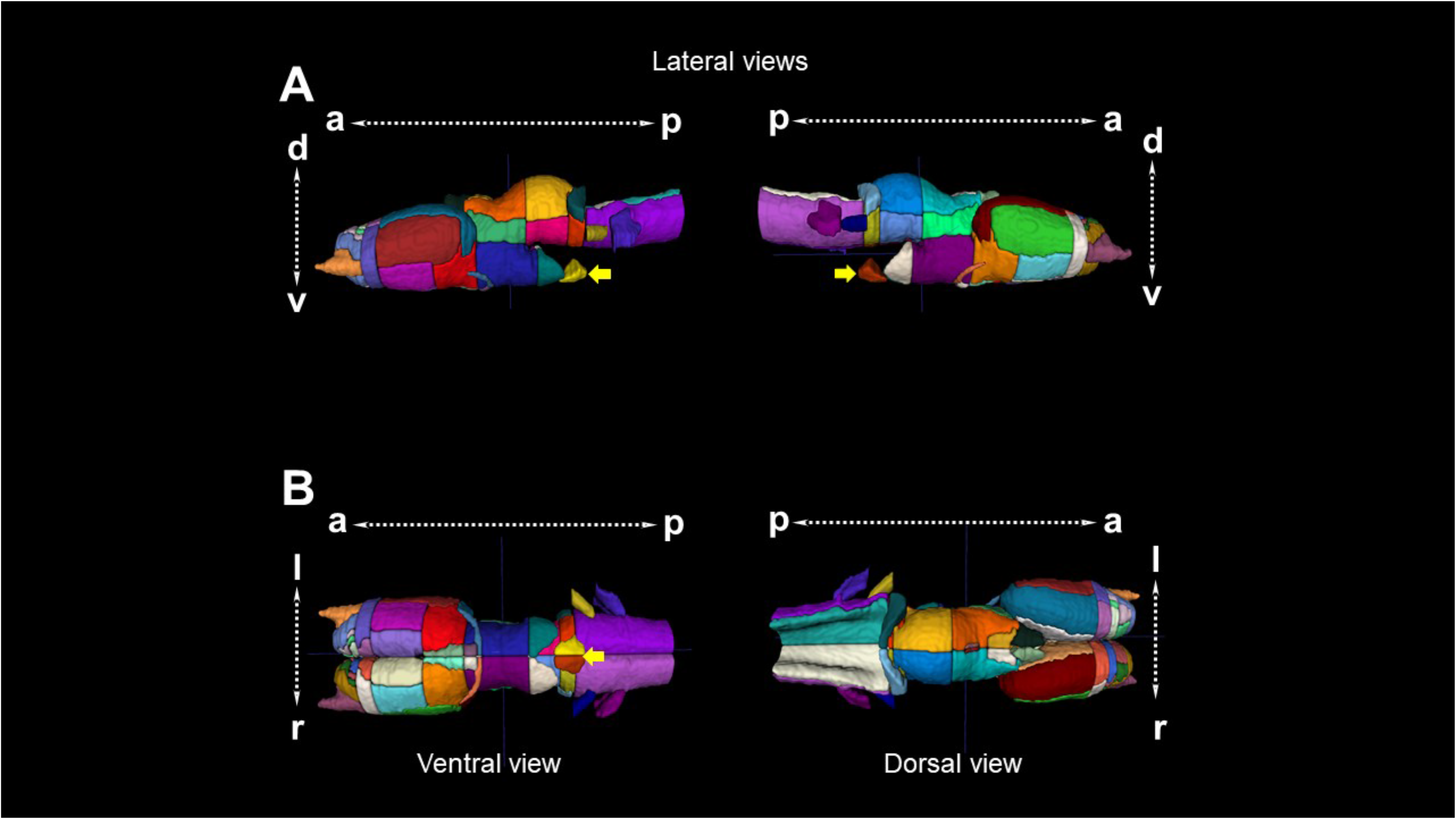
3D reconstruction of the brain of *Ambystoma mexicanum* employing MRI. A.- Lateral views from the left and right sides of the complete 3D reconstruction. B.- Ventral (left) and dorsal (right) views of the complete 3D reconstruction. Every reconstruction shows each region in a different color when manual delineation was possible. Yellow arrows denote the pituitary gland.

### Olfactory bulb

Employing previous information of the olfactory bulb structures, we are able to distinguish the olfactory nerve, the glomerular-, mitral- and granular cell layers and the anterior olfactory nucleus (**Table 1**, **Figure 2**). These structures contribute to the integration of information coming from olfactory neuroreceptors from the nasal cavity. In mammals, some of the cell structures that contribute to integrate odorant signals are well documented in terms of morphology, connectivity and function ^17^, but information in amphibians is still scarce.

**Figure 2.**
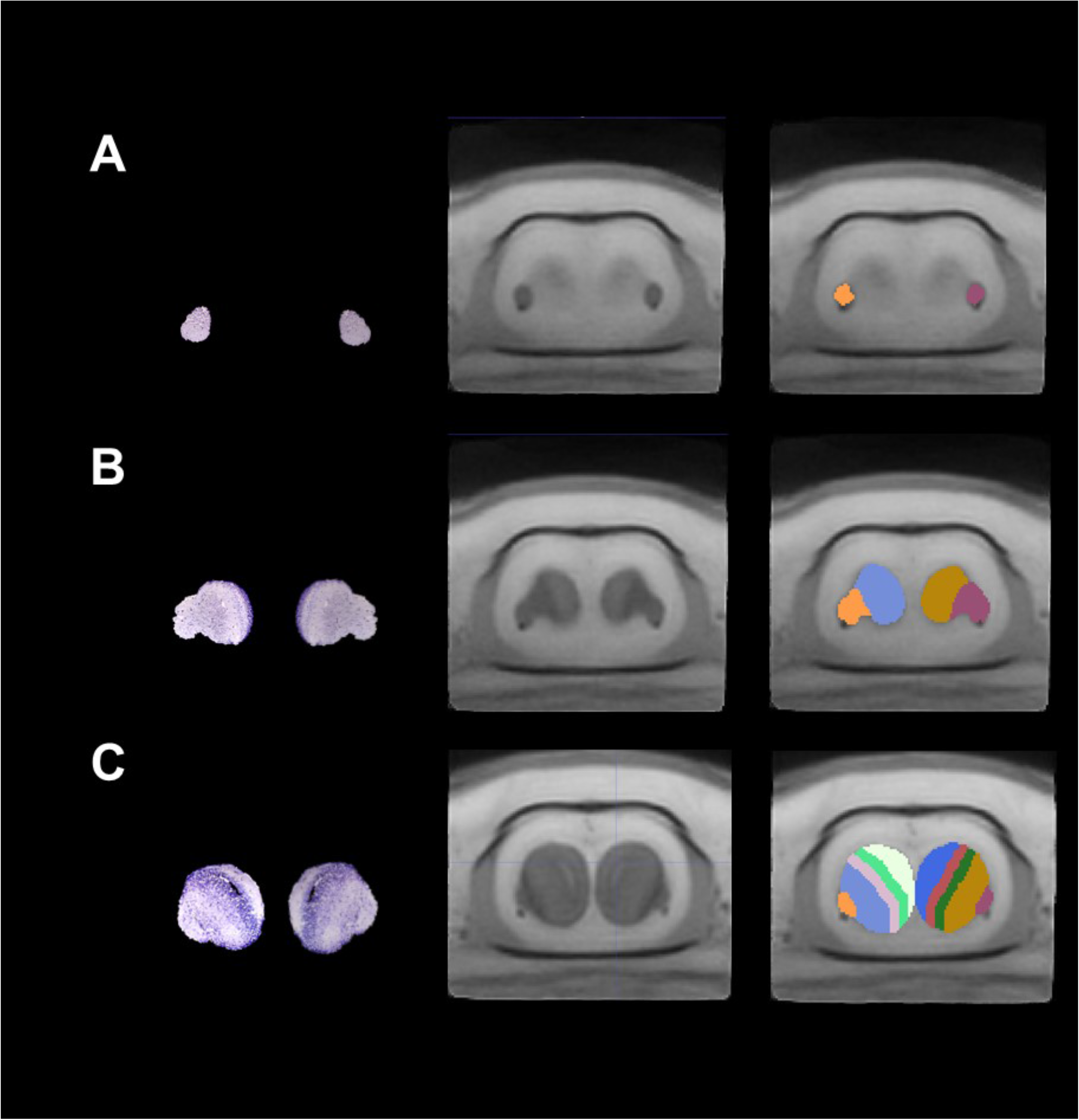
Coronal sections of the olfactory bulb of *Ambystoma mexicanum*. Columns left, central and right depict the Nissl-staining histological sections; the MRI images and the manual segmentation, respectively. (A) left and right single olfactory nerves; (B) the olfactory nerve, glomerular layer, and (C) mitral and glomerular cell layers.

### Telencephalon

This is the biggest brain structure in salamanders; the anterior part of the telencephalon contains the pallium and subpallium and the lateral ventricles. The pallium and subpallium comprise the left and right hemispheres. Axolotl pallium is interesting due to its capacity to regenerate after damage ^4^, a trait probably present in other salamanders ^7^. Because our MRI did not allow enough contrast to differentiate structures such as the dorsal pallium, lateral pallium, medial pallium and striatum and septum, we decided to further segment these regions manually using visual inspection from drawings in previous publications (**Supplementary Table 1**). The lateral ventricles (not segmented) were hyperintense in terms of contrast in our T2-weighted MRI images and are depicted in **Figure 3A**. The dorsal, lateral and medial pallium extend to the posterior part of the telencephalon but present a different morphology as compared with the anterior telencephalon structures. Moreover, at this brain location, the distance between the lateral ventricles (left and right) is reduced and even located adjacently at a more posterior region, finally forming the third ventricle (**Figure 3B)**. We segmented the amygdaloid complex, the lateral/medial forebrain bundle, the anterior preoptic nucleus, the thalamic eminence, and the pallial and anterior commissures, according to previous annotations **(Figure 3C).** Moreover, the choroid plexus, which secretes cerebrospinal fluid into the vertebrate brain ^18^ was evident using MRI, contrary to what is observed in histological analysis due to the difficulty to preserve this structure.

**Figure 3.**
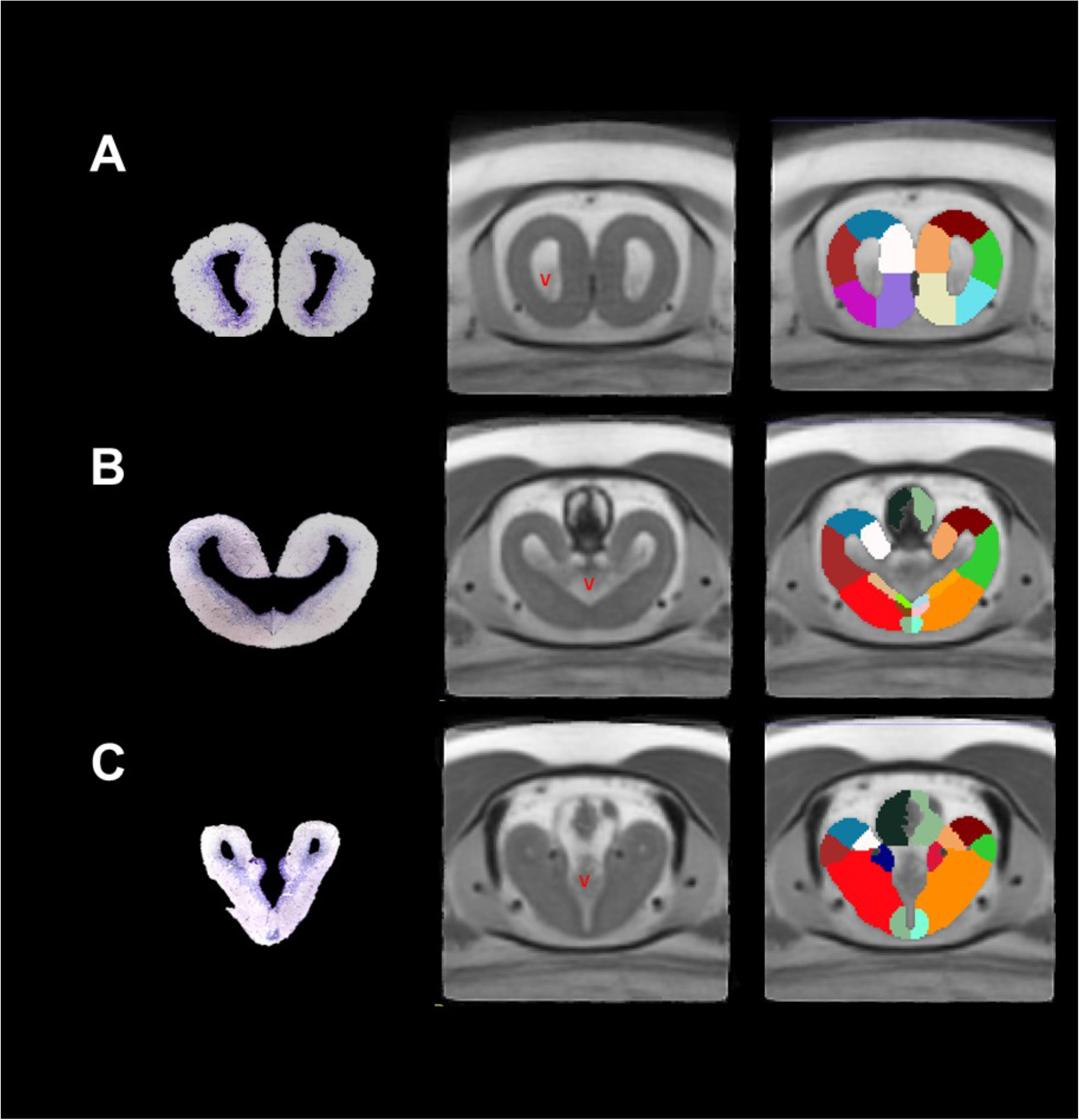
Coronal sections of the telencephalon of *Ambystoma mexicanum*. Columns left, central and right depict the Nissl-staining histological sections; the MRI images and the manual segmentation, respectively. (A) pallium (dorsal, lateral and medial pallium) and subpallium (striatum and septum); (B) choroid plexus, amygdaloid complex, pallial and anterior commissures, and (C) thalamic eminence and anterior preoptic nuclei. Ventricles (V) are indicated.

### Diencephalon

At the diencephalon level, we could identify two general structures, thalamus and dorsal hypothalamus (**Figure 4**). We segmented the thalamus into dorsal and ventral thalamus. At the top of the dorsal thalamus, we identified the habenula (**Figure 4A)** and the subcommissural organ (**Figure 4B)**. The habenula is a structure present in all vertebrates which participates in the integration of the limbic system, the basal ganglia and the sensory information ^19^. The subcommissural organ contains secretory ependymal cells located at the roof of the third ventricle ^20^. The contrast of the latter was hyperintense and centrally located with respect to the thalamus/hypothalamus. The dorsal hypothalamus includes nuclei that surround the floor of the third ventricle such as the posterior preoptic nucleus, paraventricular organ, pars dorsalis and pars ventralis hypotallami. All these regions have been recognized as neurosecretory cells which participate in neuroendocrine systems (**Figure 4A and B)**^21^. Finally, the optic chiasm was also evident in our MRI images at the floor of the dorsal hypothalamus (**Figure 4A)**. In this region, nerve fibers cross and allow binocular communication between eyes and the brain ^22^.

**Figure 4.**
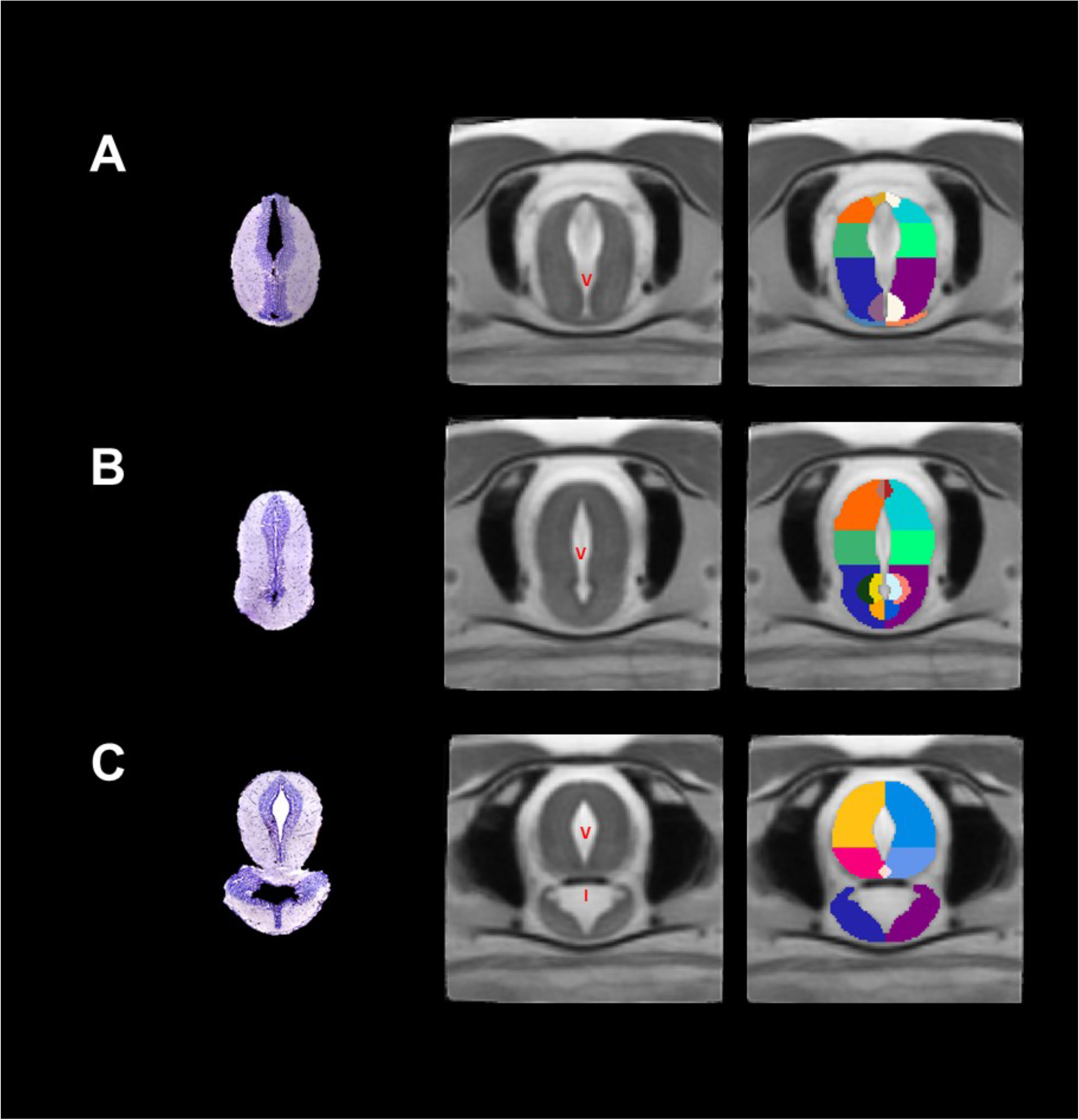
Coronal sections of the diencephalon/mesencephalon of *Ambystoma mexicanum*. Columns left, central and right depict the Nissl-staining histological sections; the MRI images and the manual segmentation, respectively. (A) dorsal and ventral thalamus, subcommissural organ, optic chiasm; (B) regions surrounding the floor of the third ventricle in the dorsal hypothalamus such as the paraventricular organ, pars dorsalis and pars ventralis hypotallami; (C) the mesencephalon, including the optic tectum, tegmentum, interpeduncularis nucleus and ventral hypothallamus. V= ventricle and I= infundibulum are indicated.

### Mesencephalon

This section was subdivided into tectum, tegmentum and ventral hypothalamus. Since contrast was insufficient to differentiate between tectum and tegmentum, we manually segmented these structures using previously published drawings (**Figure 4C**, **Supplementary Table 1**). Optic tectum is the roof of the mesencephalon, and a big region in the salamander brain; it receives some nerves from the optic chiasm and processes visual signals in vertebrates ^23^. Tegmentum is located just ventrally from the optic tectum; from this region we were able to segment the interpeduncularis nucleus, which is known to integrate information for the limbic system ^24^. At the coronal level, the ventral hypothalamus appears as the lower area of the brain, physically separated from the optic tectum/tegmentum and surrounding the infundibulum (not segmented). However, the ventral hypothalamus becomes smaller in the left and right sides at posterior levels eventually disappearing, whereas the infundibulum increases in size. The third ventricle remains in the cerebral midline and the center of the optic tectum/tegmentum.

### Rhombencephalon/pituitary

This is the most posterior area of the brain and contains regions such as the cerebellum and medulla oblongata. Cerebellum in amphibians is small in size with respect with other vertebrates ^25^, a characteristic inversely correlated to the big genome of these species ^26^. As previously reported, we detected a small cerebellum in our MRI images (**Figure 5A**). Medulla oblongata was identified as a single structure in the anterior rhombencephalon, but we were able to segment gray from white matter in the posterior rhombencephalon, according to differences in contrast in our MRI images and previous reports (**Figure 5B**, **Supplementary Table 1**). Moreover, we also recognized some cranial nerves such as nervous trigeminus (V), nervous lateralis anterior and nervus octavus (VIII) (**Figure 5B and C)**. Finally, we identified the pituitary gland, an endocrine organ which releases hormones in response to some peptides coming from hypothalamus ^27,28^. In our coronal sections, the pituitary gland appears at the lower part of the brain as the last visible signal of the ventral hypothalamus, increasing in size as it projects posteriorly to finally decrease and disappear (**Figure 5A**)

**Figure 5.**
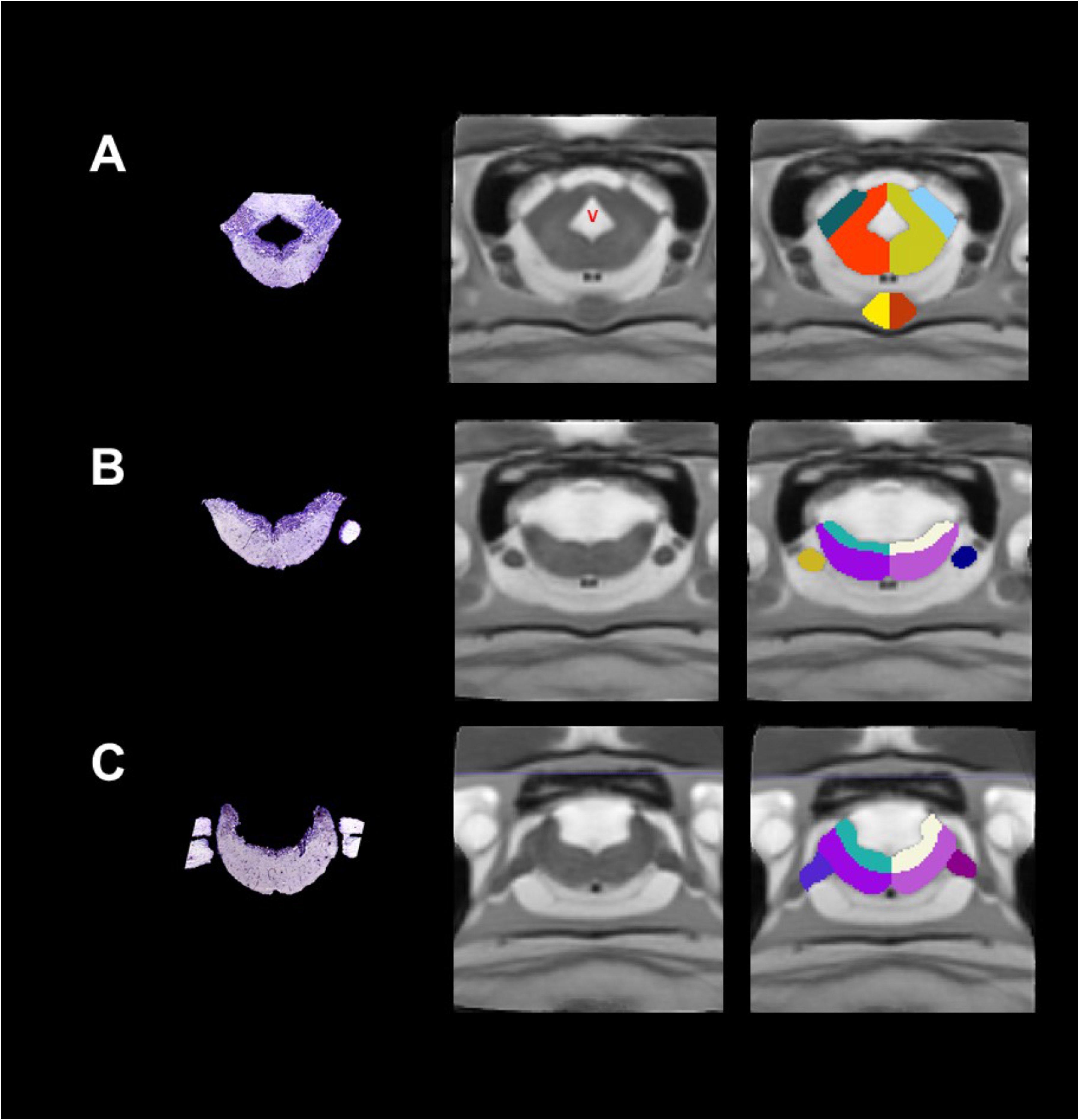
Coronal sections of the medulla oblongata and pituitary gland of *Ambystoma mexicanum*. Columns left, central and right depict the Nissl-staining histological sections; the MRI images and the manual segmentation, respectively. (A) the rhombencephalon, where the cerebellum, the medulla oblongata and the pituitary gland at the lower part of the brain are indicated; (B) white and gray matter from medulla oblongata and cranial nerves such as the trigerminus nerve, and, and (C) nervous lateralis anterior and nervus octavus.

### Myelin rich regions in the axolotl CNS

Myelin is an innovation of vertebrate CNS ^29,30^ which allows saltatory propagation of action potentials. Patterns of myelin rich regions are well-documented in mammals ^31–33^ but in non-mammalian vertebrates, particularly in amphibians, the information is scarce. In an approach to identify myelin in the axolotl CNS, and due to its contribution to T2-weighted contrast in MRI, we performed specific myelin staining using the Black Gold II reagent in sagittal and coronal sections. Myelin-rich regions were evident in sagittal sections of the medulla oblongata, cerebellum, optic tectum and tegmentum; however, no myelin staining was observed in the anterior part of the brain, *i.e.*, in the olfactory bulb and telencephalon (**Figure 6**). These results were confirmed using coronal sections, in which we are able to detect myelin in the lateral/medial forebrain blundle/amygdaloid complex (**Figure 7A**), the ventral thalamus (**Figure 7B**) and the optic tectum/tegmentum (**Figure 7C**). Both, sagittal and coronal sections showed that medulla oblongata is a myelin rich region, particularly in the white matter, confirming the specificity of the myelin staining method (**Figure 7D**). We were not able to detect myelin in the anterior part of the axolotl brain. This discrepancy could have different explanations. In the developing mammal, the myelination process starts in the spinal cord and gradually covers the posterior part of the CNS postnatally ^34,35^. It is possible that even if the juvenile stage of the axolotl brain is not fully myelinated, the adult brain could contain myelin in the anterior part of the brain. Another explanation could be that the axolotl lost the capacity to myelinate axons in the anterior part of the brain as result of neoteny and/or other unidentified physiological events. In this context, all amphibians go through dramatic transformations that occur in the transition from a pre- to a post metamorphic stage; however, axolotl can retain juvenile features throughout adulthood (neoteny/paedomorphism), resulting in adult (reproducing) individuals that maintain juvenile (larval) traits). Lastly, amphibian myelin composition might not be detected by Black Gold II. Even if this reagent is indeed myelin specific, amphibians have shown to present different compositions of myelin in terms of lipid content ^36,37^. Therefore, it is plausible that axolotl could have a myelin that is not efficiently stained with Black Gold II in the anterior brain, at least at the stage of development that was analyzed in the present study. To confirm this last hypothesis, other stages of axolotl development and/or other methodologies to detect myelin could be tested.

**Figure 6.**
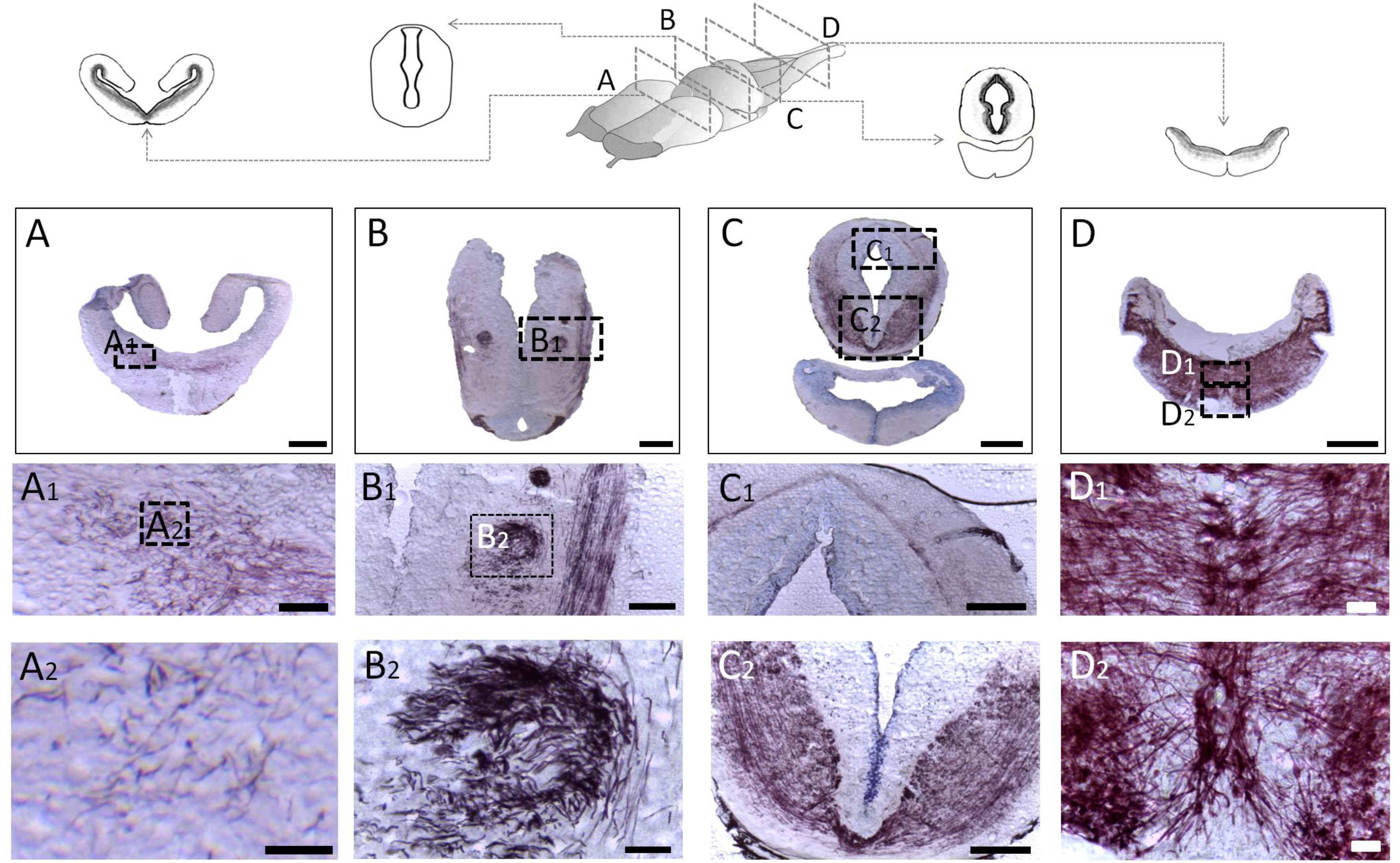
MRI and BGII techniques in the axolotl brain. (A, C) sagittal sections of in-vivo T2-weighted images and myelin staining with BGII; (B, D) the approximate myelin localization is indicated in the diagram on the left. Insets in A and C show diagrams of the corresponding sagittal plane. In the histological micrographs of BGII staining, E and G are amplifications of B and D, while F and H are amplifications of E and G, respectively. Bars in B, D indicate 1mm, in E, G they indicate 100um, while in F and H they indicate 20um.

**Figure 7.**
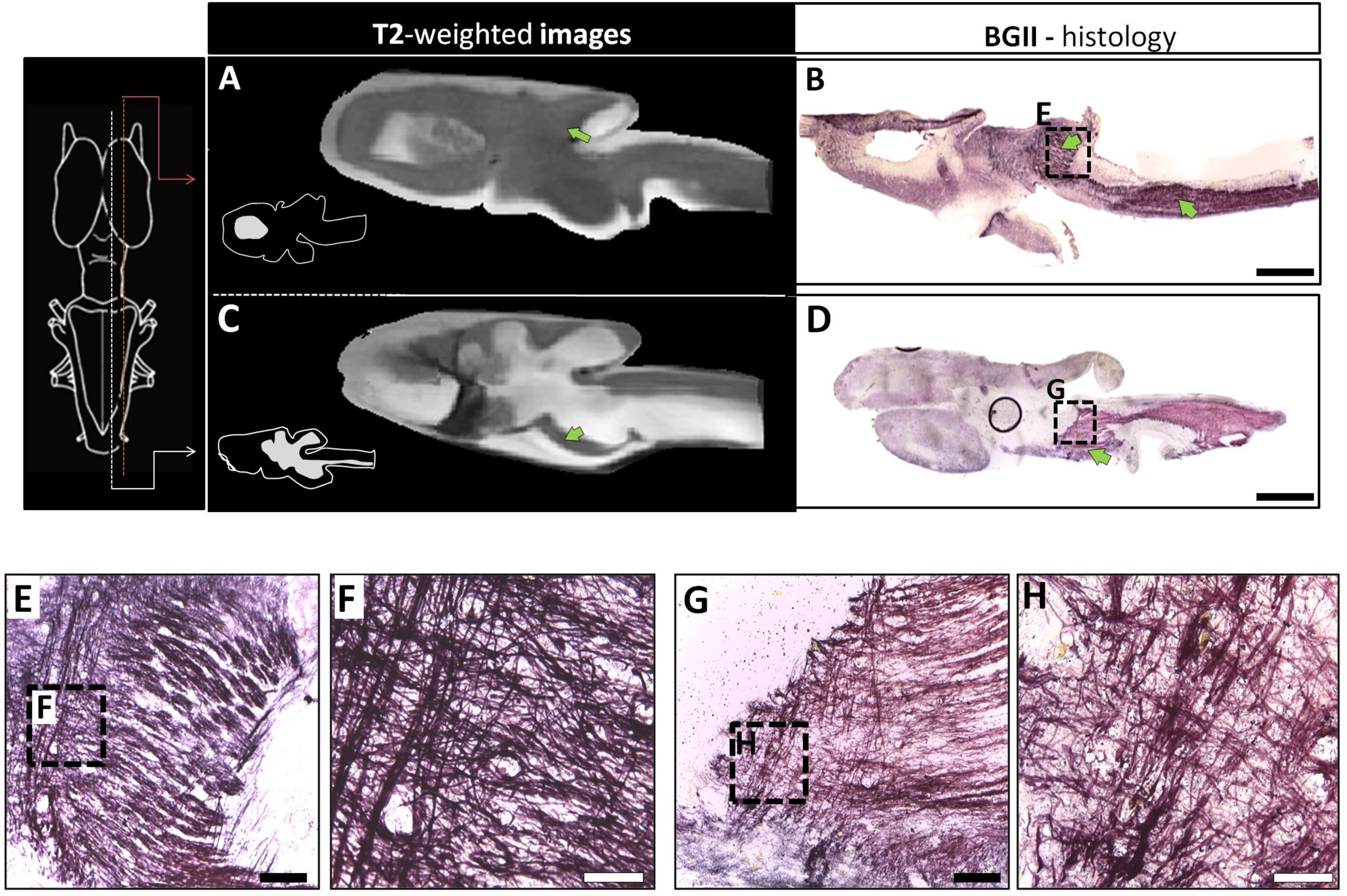
Myelination in the axolotl. Different coronal sections of the axolotl brain. (A) lateral/medial forebrain bundle/amygdaloid complex; (B) the dorsal thalamus; (C) the optic tectum/tegmentum, and (D) medulla oblongata. Below are amplifications for each section. Note the myelination gradient in the anteroposterior direction. The white matter of medulla oblongata is the structure with the greatest presence of myelin. The bars in the micrographs indicate 500 μm in A-D, 100 μm in A1, B1, C1, C2, and 20 μm in A2, B2, D1 and D2.

## Conclusions

The brain of vertebrates has been studied with different techniques, allowing researchers to visualize different aspects of its anatomy, according to the methodology employed. MRI of the axolotl brain allowed us to provide a 3D reconstruction of an amphibian brain and pituitary gland for the first time. In contrast with the results from histological sections, we were able to visualize native morphology of the brain structures and to transform this data in a 3D volume. Additionally, this is the first description of axolotl brain myelin distribution. Myelin rich regions were observed in the posterior, but not the anterior brain, a finding that deserves further attention. Overall, this work will provide a useful tool to explore new research avenues for the better understanding of this interesting endemic paedomorphic species, which is an important model in different research areas, including nervous tissue regeneration, and that is currently listed as critically endangered by the International Union for Conservation of Nature.

## Materials and Methods

### Animals

Juvenile axolotls were kindly donated by Marco Terrones (Axolkali). All axolotls were maintained and handled in accordance with protocols approved by the Ethics for Research Committee of the Instituto de Neurobiología at the Universidad Nacional Autónoma de México (UNAM). Animals of around 3 months after hatching and weighing between 9-12 g were kept at our local housing for at least 20 days prior to imaging at 18° C in 14/10 light/dark cycles. After habituation, animals were anesthetized by immersion with 0.4 % tricain for 10-13 minutes.

### MRI acquisition

MRI was performed on 20 animals at the National Laboratory for MRI, using a 7 Tesla Bruker Biospec 70/16 scanner and a Helium-cooled two-channel rat-head coil (Bruker Cryoprobe). Once anesthetized, axolotls were placed in a plastic container and introduced in the scanner (**Supplementary Figure 2**). A field map was first obtained and used for shimming of the main magnetic field. Next, images were acquired using a three-dimensionally encoded balanced steady-state gradient echo (True FISP) sequence with the following parameters: TR=4.4 ms; TE=2.2 ms; flip angle=30°; NEX=3; FOV=20.48×15×10.24 mm^3^ and matrix=256×188×128, yielding isometric voxel dimensions of 80 μm^3^; scan time= 5 min 50 sec. We also acquired diffusion-weighted images with a scan time of 11 min 44 sec (not reported herein). Total scan time was 17 min 34 sec.

### MRI template construction

Individual image volumes were visually inspected for quality. Out of n = 20 images, n = 6 were rejected due to poor quality, with a final sample size of n = 14 for template construction. Images were converted from Bruker format to NIFTI using the software Bruker2nifti ^38^, and then from NIFTI to MINC using nii2mnc (https://github.com/BIC-MNI/minc-toolkit-v2). Several steps were followed to reach the final template construction. STEP1: each image was preprocessed using the following steps and commands; 1) cleaned header and center image to coordinate 0,0,0 (https://github.com/CoBrALab/minctoolkitextras/blob/master/clean_and_center_minc.pl); 2) 90 degrees rotation of the *y axis* using volrot; 3) *y axis* volume flip using volflip; and 4) N4 bias field correction using an Otsu mask ^39^. STEP 2: we constructed a first template using the antsMultivariateTemplateConstruction2.sh script (https://github.com/ANTsX/ANTs/blob/master/Scripts/antsMultivariateTemplateConstruction2.sh) ^40^. STEP 3: an initial brain mask was manually defined from the first template. STEP 4: fine tuning of the N4 correction and the first template; 1) inverse registration of the template brain mask to the native space for each subject; 2) reduce field of view to near brain (crop) using ExtractRegionFromImageByMask; 3) re-run the N4 bias field correction with the improved individual mask; 4) re-center the images; 5) rotation of the first template to a precise alignment with the axes. STEP 5: construction of the second template using the raw files and the first template as target. STEP 6: resampled second template from 0.08 × 0.08 × 0.08 mm to 0.06 × 0.06 × 0.06 mm. STEP 7: construction of the third template using the second template for warping. STEP 8: resampled final template from 0.06 × 0.06 × 0.06 mm to final resolution of 0.04 × 0.04 × 0.04 mm. STEP 9: construction of the final template using the third template for warping. The process is shown in **Supplementary Figure 3**.

### MRI segmentation

Neuroanatomical segmentation was done manually drawing regions of interest (ROIs) using the ITK-SNAP (version 3.8.0) ^41^ on the final template (0.04 × 0.04 × 0.04 mm), based on the available histological anatomic annotations. We determined and localized 80 ROIs identified in previous histological atlases (see **Supplementary Table 1** for the reference of every region). The ROIs were drawn while examining all three stereotaxic planes to reduce inconsistencies in delineation across slices. Labels were hierarchically constructed to simplify MRI analysis in higher resolutions. There are 3 levels that describe larger regions: 1) ROI; 2) embryological origin of the neural tube (olfactory bulb, telencephalon, diencephalon, mesencephalon, endocrine and rhombencephalon), and 3) hemispheres (right and left). Each ROI has a defined abbreviation. The atlas is openly available through Zenodo (DOI: 10.5281/zenodo.4311937).

### Histological staining

For histological examination, brains were fixed in PFA (4 %) and cryopreserved in sucrose (30 %), then coronal or sagittal (40 μm) cryosections were cut and mounted on electrocharged slides and stored at −20 °C until staining with the Black Gold II compound (EMD Millipore Corp., Billerica, MA, USA), using a modified protocol described by ^42^. Briefly, tissue sections were rehydrated in distilled water for about 2 min at RT. Then, the slides were incubated in 0.3 % Black Gold II solution, dissolved in 0.9 % saline vehicle (NaCl), and heated at 60-65 °C for at least 40 min until the staining was complete. Next, the slides were rinsed for 2 min in Phosphate-Buffered Saline (PBS, RT), transferred to a 1 % sodium thiosulfate solution for 3 min at 60–65 °C, and rinsed two or three times with distilled water. Sections stained with Black Gold II were counterstained with Nissl. Additional alternate slides were stained only with the Nissl protocol. For this, slides were transferred to a solution of 0.1–0.2 % cresyl violet (blue; EMD Millipore Corp., Billerica, MA, USA) in 0.1 % acetic acid for 5 min and then rinsed two times with distilled water. After, the tissue was dehydrated using sequential graduated alcohol solutions (50, 70 and 96 %) and immersed in xylene for 1 min. The slides were then coverslipped with Entellan mounting medium.

## Acknowledgements

The authors would like to thank the technical assistance of M. C. Patricia Villalobos, Guadalupe Yasmín Hernández Linares, M.C. Leopoldo González Santos, and Dr. Ericka A. de los Ríos. This work received support from Luis A. Aguilar (Laboratorio Nacional de Visualización Científica Avanzada), and the Laboratorio Nacional de Imagenología por Resonancia Magnética. We would also like to thank Gabriel A. Devenyi for his support and guidance. This work was supported by a grant from PAPIIT IN204920.

## Author contributions

IL: Conceptualization, methodology, investigation, writing - review & editing, formal analysis, resources, writing - original draft, writing - review & editing, visualization.

ACM: Methodology, investigation, writing - review & editing, visualization.

LC: Conceptualization, writing - review & editing, supervision

JJOR: Methodology, investigation.

EAGV: Conceptualization, methodology, software, formal analysis, resources, writing - original draft, writing - review & editing, visualization.

AO: Conceptualization, project administration, supervision, funding acquisition, writing - original draft, writing - review & editing, visualization

