## Supplementary material for "MRI- and histologically derived neuroanatomical atlas of the *Ambystoma mexicanum* (axolotl)"

**Supplementary Figure 1.** Brain volume per hemisphere.

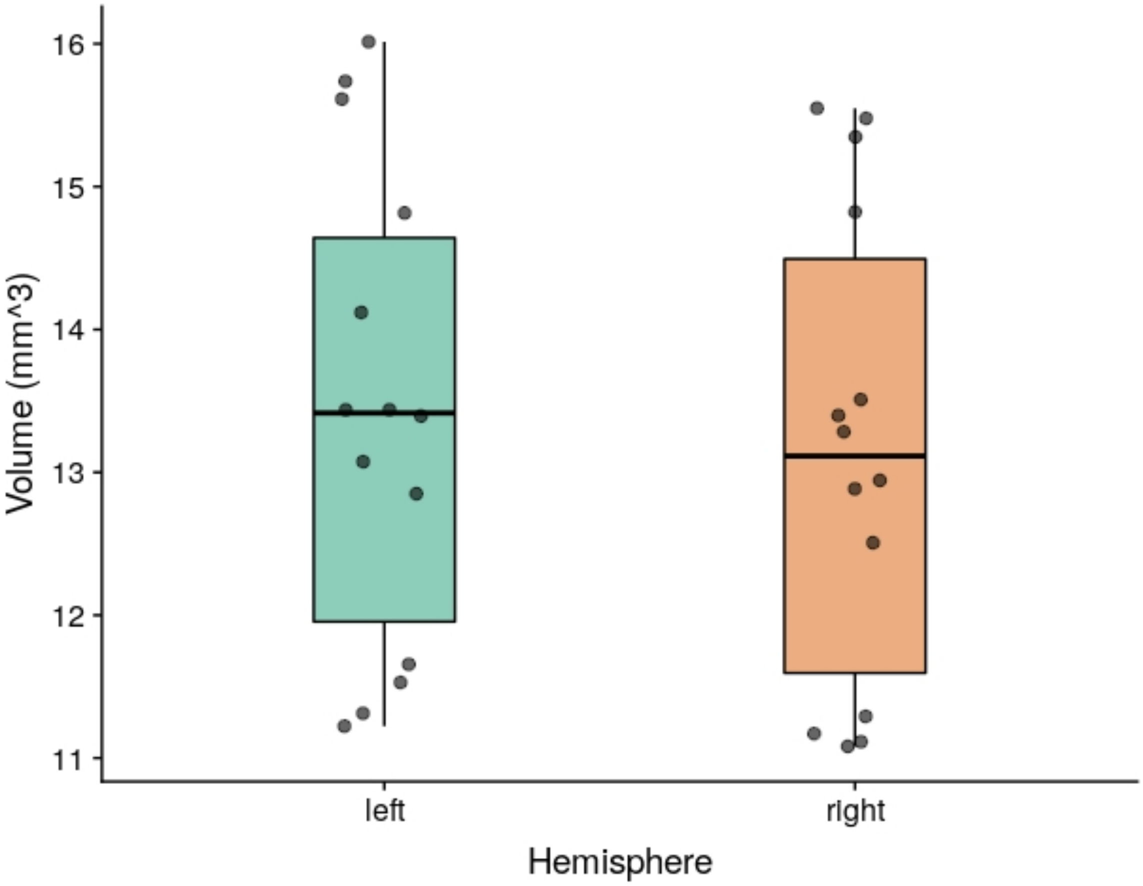


Average volume ($mm^3$) of left and right hemispheres from 14 juvenile axolotls.

**Supplementary Figure 2.** MRI data aquisition

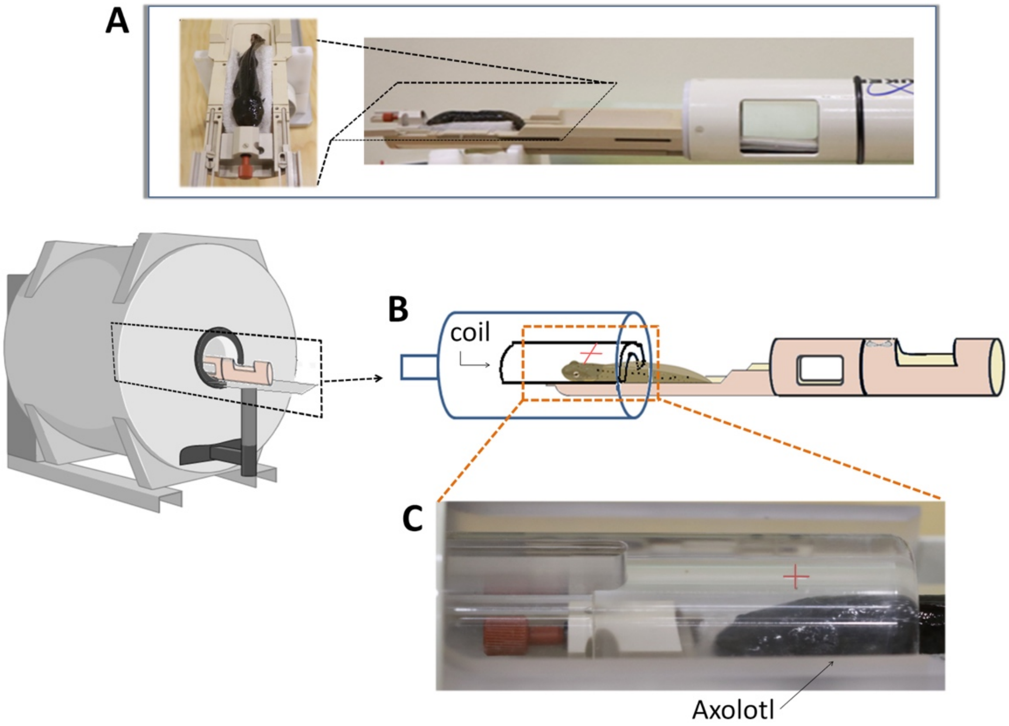


*Once anesthetized, the axolotl is placed on a plastic stretcher over a holder, in (A) the axolotl is observed in a top view (left) and from a lateral view (right). Then, it is inserted into the scanner, in such a way that the center of coil (Helium-cooled two-channel rat-head coil: Bruker Cryopobe) is approximately 1 cm from the eyes towards the caudal part (B). In (C) the placement of the axolotl in the coil-simulator is shown, the red mark indicates the center of the coil.*

**Supplementary Figure 3.** Pipeline for template construction.

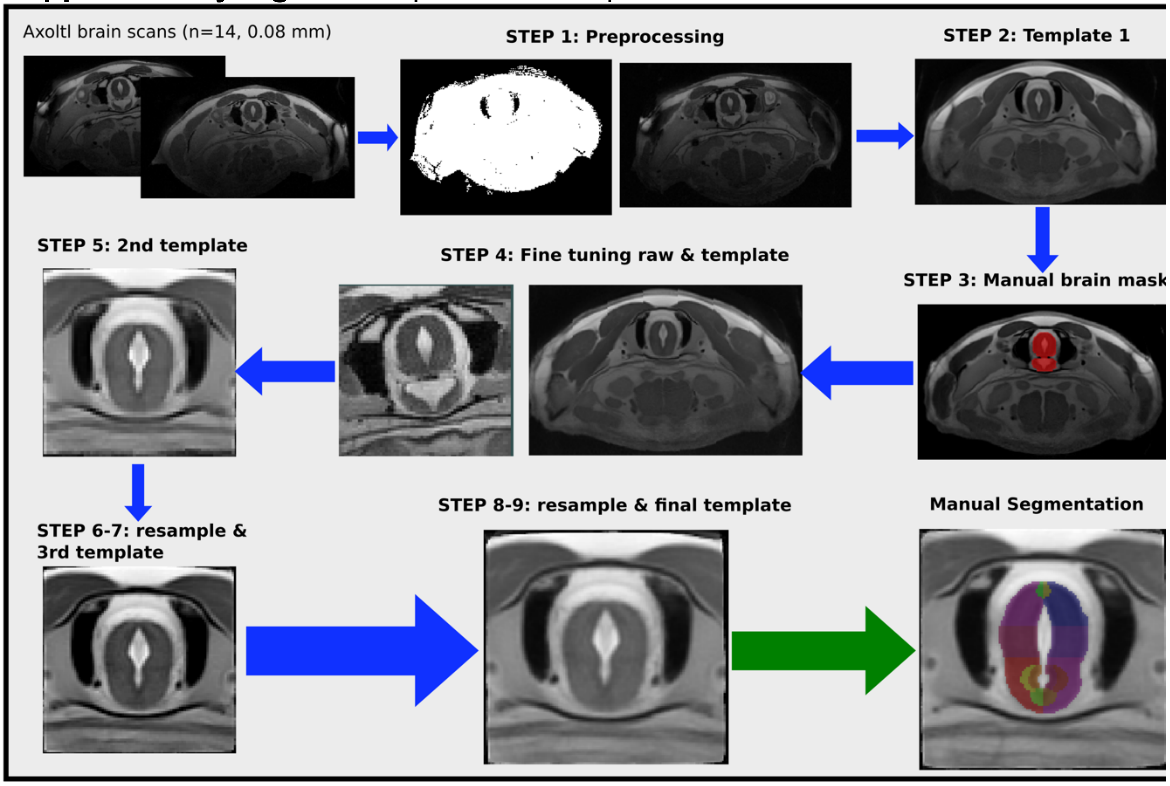


*Flow chart of the process used to construct the template used for segmentation. Details about the steps in the Methods Section.*
