## Supplementary Table 1 for "MRI- and histologically derived neuroanatomical atlas of the *Ambystoma mexicanum* (axolotl)"

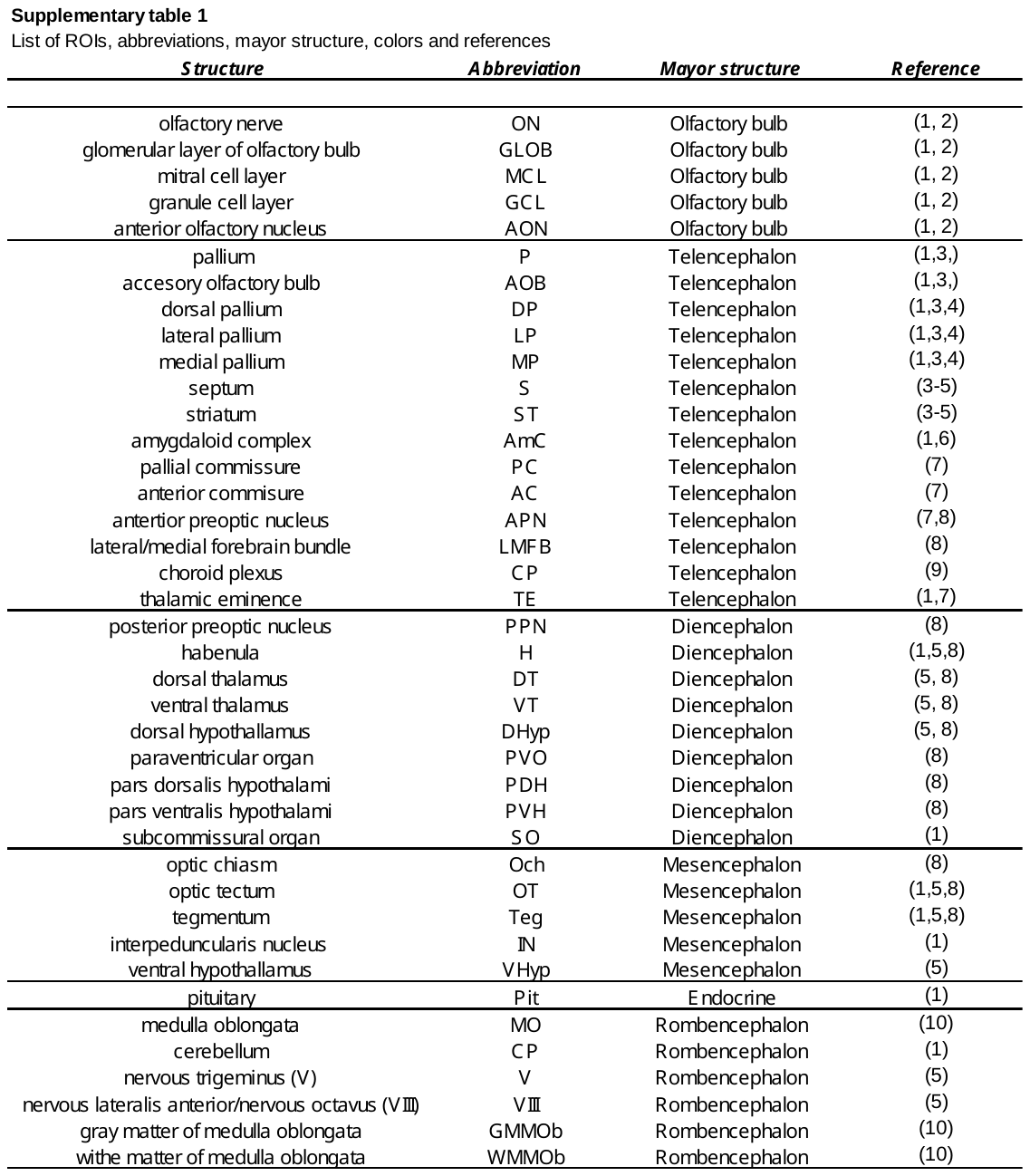
